# Haplotype and Repeat Separation in Long Reads

**DOI:** 10.1101/145474

**Authors:** German Tischler

## Abstract

Resolving the correct structure and succession of highly similar sequence stretches is one of the main open problems in genome assembly. For non haploid genomes this includes determining the sequences of the different haplotypes. For all but the smallest genomes it also involves separating different repeat instances. In this paper we discuss methods for resolving such problems in third generation long reads by classifying alignments between long reads according to whether they represent true or false read overlaps. The main problem in this context is the high error rate found in such reads, which greatly exceeds the amount of difference between the similar regions we want to separate. Our methods can separate read classes stemming from regions with as little as 1% difference.

## 1 Scientific Background

Third generation sequencing reads like those produced by Pacific BioSciences (Pac-BIO) and Oxford Nanopore Technologies (ONT) sequencers are very long in comparison with the ones produced by second generation sequencers. The average read length for PacBIO is often 10k base pairs (bp) and for ONT 7-9kbp have been reported. For PacBIO more than half of the sequenced bases can be in reads of length 20kbp and above. This is much longer than the reads produced by second generation sequencers, which often yield reads as short as 150bp, so third generation sequencers allow a much better repeat resolution for assembly because more repeats are spanned by single reads. This increased read length however comes at the price of a much higher average base error rate (about 13% for PacBIO and even higher for ONT). This poses major algorithmic challenges in the areas of sequence alignment, comparison and signal detection. Read versus read comparison (see e.g. [1]) operates at a correlation of 70% and less. This makes it very hard to detect small differences between regions reads were sampled from. Reads stemming from sufficiently similar regions in an underlying genome, like instances of a repeat or different haplotypes, will often align within the parameters used, as the difference between the two sources is small in comparison with the read error rate. Being able to segregate read alignments into classes according to whether or not an alignment between a read pair designates a real overlap in the underlying genome to the degree possible is however important for multiple applications like genome assembly and variant detection. In genome assembly for instance the quality of any consensus sequence produced rises and falls with the ability to select the correct reads as input (cf. [2]). Linking up different haplotypes during the assembly of a non haploid organism results in patchwork like output, in particular an assembly process yielding output contigs which are in this way not contained in the genome to be reconstructed. Haplotype assembly designates the problem of separating reads into haplotype classes by first mapping them to a given reference sequence and then splitting the reads into groups using the information obtained. Several papers have presented methods for haplotype assembly (or read phasing) in the diploid setting (cf. [3, 4, 5, 6, 7, 8]). Canu (see [9]) performs repeat separation using a sequence of error correction, residual error estimation and classification of the error corrected reads. The authors report being able to separate repeat instances with 3% difference and above.

## 2 Materials and Methods

In this paper we discuss methods for repeat and haplotype separation in long reads. We consider the setting of de novo assembly, in particular we do not presume or require the existence of a reference sequence or known variation sites. Instead of read to reference alignments we use read to read consensus alignments, i.e. we align reads to error corrected reads. In addition we do not limit our attention to a scenario requiring there to be at most two versions of a sequence, like it is the case for the haplotype assembly of diploid genomes.

We consider two basic principles for splitting a set of reads. The first one is based on the trivial observation that reads in the same class should agree on most positions, particularly including those for which a very rough analysis shows a potential for disagreement in the read set. As the reads we consider are not error free we cannot expect the reads inside one class to agree on all positions. This approach has it’s merits when the number of versions a sequence appears in is low but as we will see below, it becomes unsuitable as the number of versions grows. The second principle is based on observing sets of reads (more or less) consistently disagreeing on certain positions. This scales to higher version numbers but is computationally much more expensive.

### 2.1 Preliminaries

Let *G* = {*S*_1_, *S*_2_,…, *S_k_*} denote a genome containing sequences *S_i_* for *i* = 1,…, *k*, i.e. strings over the alphabet Σ = {*A,C,G,T*}. Further let *R* = {*R*_1_, *R*_2_,… *R*_r_} be a set of reads sampled from *G* (randomly of the forward and reverse complement strand) such that the strings in *R* have length *L* on average and the error rate (errors per length on *G*) between the reads and the intervals on *G* they were drawn from is *p*_*e*_ on average. For PacBIO the length distribution in R would follow a log normal distribution with average length 10kb and *p*_*e*_ would be in the order of 0.13. We denote a local alignment between sequences *U*_*i*_ and *U_j_* by a tuple (*i,j,ib,ie,jb,je,c*) where ib and ie mark the start and end of the alignment on *U*_*i*_, *jb* and *je* the start and end on *U*_*j*_ and *c* is a Boolean value marking whether *U*_*j*_ or the reverse complement of *U*_*j*_ was used (*c* = true for reverse complement). In practice long reads often contain stretches of very low quality, so even for two reads sharing a true overlap we find a sequence of local alignments instead of a single suffix/prefix or containment type alignment. Our methods can easily be generalised to this case, however for the sake of simplicity of exposition we assume that alignments between reads are contiguous below. Let *A* denote the set of all (local) alignments between pairs of reads in *R* s.t. the correlation between the two reads inside the alignment is at least 1 — *2p*_*e*_ and the alignment covers at least *l* bases on both reads involved for some length *l*. In practice we commonly use *l* = 1*k* for third generation long reads. For a given read *R*_*k*_ we call the subset of *A* s.t. the first component of the tuples is *k* the alignment pile for *R*_*k*_. If *G* represents a non haploid genome or contains sufficiently long repeating regions then not all the alignments in *A* may refer to true read overlaps on *G*. We can use an alignment pile of a read or a subset thereof (for instance by choosing the top *k* best aligning other reads for some *k*) to compute a preliminary consensus or error corrected version of the read, e.g. using the algorithm proposed in [2]. We denote a preliminary consensus obtained for a read *R*_*i*_ in this way by 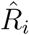. The alignment pile for *R*_*i*_ can be transformed into an alignment pile for 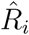 by aligning the reads in the pile for *R*_*i*_ to 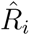 while taking the positions of the original alignments on *R*_*i*_ into account and transforming those to positions on 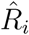 using an alignment of *R*_*i*_ and 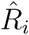. In addition to the original alignments in the pile of *R*_*i*_ we also insert an alignment between 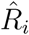 and *R*_*i*_ into the pile of 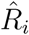. An alignment pile for some 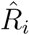 can be transformed into a matrix where the columns represent positions on or before the bases of 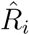 (before for base insertions into 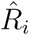) and the rows are the reads in the alignment pile for 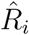. An alignment (*k,j,kb,ke,jb,je,c*) between 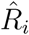 and a read *j* is active from base *kb* to ke on *k*. The cells of a matrix row corresponding to read *R*_*j*_ are set as follows. Columns the respective alignment is inactive on remain empty. In the active region of the alignment a cell is filled with the base from *R_j_* if the alignment features a match, mismatch or insertion operation for the respective position and a dash (—) otherwise. As a convention we always have the alignment between 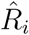 and *R*_*k*_ as the first row of the matrix. Table 1 shows an example. The excerpt shows positions 100 to 102 on the read. Some bases have been inserted before position 102, which is marked by the position identifier (102, –1). The alignment corresponding to the second row ends at position 101, the one for the last row starts at position 101.

**Table 1.** Excerpt from a matrix given by an alignment pile

|  |  |  |  |
| --- | --- | --- | --- |
| (100,0) | (101,0) | (102,-1) | (102,0) |
| A | C | A | T |
| A | C |  |  |
| A | T | A | T |
|  | C | — | G |

### 2.2 Agreement Based Splitting

Let *d* denote the average sequencing depth of the read set *R*. We assume the arrival rate of reads on the genome follows a Poisson distribution with mean *d*, i.e. we have a probability of 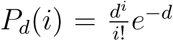 to see a depth of i at a given position. The probability to see *d’* correctly sequenced bases for any position is thus

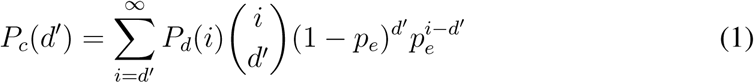

We want to detect variation sites inside a given read *R*_*i*_. One very simple way to do this is to scan the matrix constructed for 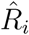 for columns in which more than one symbol appears with a frequency above a given threshold. Assuming the alignments used to construct the matrix are suitable we would see such a variation with probability 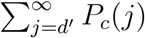 if we chose a threshold of *d’*. For *d* = 20 and *p*_*e*_ = 0.15 we obtain *d’* = 8 if we ask the probability to be at least 99%, i.e. we are 99% sure not to miss a relevant site if we look for columns containing at least two symbols with 8 or more instances. There is however the chance of calling variation sites because of unsuitable alignments in the pile for 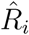 or a sufficiently high number of wrongly sequenced bases (this is a problem especially in the presence of a high number of sequence versions as this increases the total number of reads involved in the pile). Consider a given position q in the genome G and two reads *R*_*i*_ and *R*_*j*_ covering this position. Then we have a probability of (1 — *p*_*e*_)^2^ for having the base at position *q* sequenced correctly in both *R*_*i*_ and *R*_*j*_. Let *A*_*ij*_ = (*i,j,ib,ie,jb,je,c*) denote an alignment between read *R*_*i*_ and *R*_*j*_ and assume we called *n* variants on *R*_*i*_ inside the index interval [*ib,ie*]. If *R*_*i*_ and *R*_*j*_ overlap as designated by *A_ij_* in the underlying genome, then we have a probability of

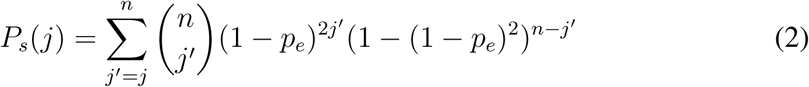

to see *R*_*i*_ and *R*_*j*_ agree on at least *j* of the *n* disagreement points in the matrix for the alignment pile of 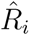. If there are two underlying versions, e.g. a repeat with two copies or haplotypes in a diploid genome, then we would expect to see reads coming from different versions disagree on most of the variant locations. In this case we have a strong signal for separating the two versions. It becomes weaker in the presence of more versions when some of the versions agree with others in a large fraction of the variant locations. In this case we cannot reliably tell the difference between two reads stemming from different versions with a relatively low number of sequencing errors and two reads stemming from the same version but agreeing on a lower number of variant locations due to a higher number of sequencing errors. For experiments we choose the number *m* of disagreement points two reads need to agree on so we consider them as from the same class as the smallest number s.t. *P*_*s*_*(m) ≥* 0.995.

### 2.3 Disagreement Based Splitting

One of the main problems with agreement based splitting is suboptimal performance when reads from different classes agree on a large number of the detected variant locations. Splitting based on the differences between genomic regions does not suffer from this effect. Every attempt via directly comparing two long reads is however bound to fail as the high sequencing error rate drowns any slight difference between the two underlying real sequences. At a single base error rate of *p*_*e*_ = 15% the probability to see a correct pair of corresponding bases in two reads is (1 — *p_e_*)^2^ = 72.25%, i.e. 27.75% of the pairs are wrong and most of these wrong pairs lead to a false disagreement between reads which should agree. When we compare bases for discovering disagreements between reads, we need to make reasonably sure that the bases compared are correct representations of their class for a given position. Consider some position on *k* reads stemming from the same class. Then we have a probability of 1 — *p_e_^k^* to see the correct base in at least one of these *k* reads. For *k* = 2 and *p*_*e*_ = 0.15 we have a probability of 97.75%, still a probability of more than 2% for all the bases to be wrong, for *k* = 3 we reach 99.6625%. In consequence, if three reads from the same class agree on a base, then this is most likely a correctly reported base. We use this observation by instead of comparing single read bases to single read bases comparing three tuples of bases to three tuples of bases. Given a read *R*_*j*_ we first build the matrix corresponding to the alignment pile of 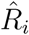. We then scan the matrix column for column. In each column *c* we extract all 6 tuples (*r*_1_, *r*_2_, *r*_3_, *r*_4_, *r*_5_, *r*_6_) s.t. *r*_*i*_ for 1, 2,…, 6 are row identifiers marking non empty cells in column c, the cells for row *r*_1_, *r*_2_ and *r*_3_ all contain the same symbol *a*, the cells for row *r*_4_, *r*_5_ and *r*_6_ all contain the same symbol *b*, *a* ≠ *b*, 1 = *r*_1_ < *r*_2_ < *r*_3_ and *r*_4_ < *r*_5_ < *r*_6_. Remember row 1 in the matrix refers to the alignment between 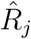 and *R*_*j*_. There are 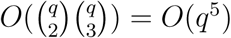 distinct such tuples in the worst case if *q* is the maximum number of active alignments in any column of the matrix. For each distinct tuple *T* we count the number *Y (T*) of times it appears summed up over all columns. The support *Z(T*) of a tuple *T* is the intersection of the active intervals of the alignments it is based on. If we want to split read sets down to a difference rate of *δ*, then we expect Δ = *δ*|*Z*(*T*)| differences to exist inside *Z*(*T*). Assuming suitable alignments comprising 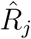’s matrix, the probability to see each single of these differences is *p*_*6*_ = (1 — *p_e_*)^6^ which is about 37.7% for *p*_*e*_ = 15%. The probability to see at least *m* of these differences is 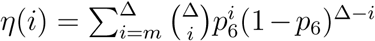. We choose the smallest *i* s.t. *η(i*) is at least 99.5% as a threshold. For each *i* we count the number *H*(*i*) of tuples satisfying their threshold in which *i* appears as *r*_4_, *r*_5_ or *r*_6_. Given *P*_*d*_ (Poisson distribution) as defined above we can determine a depth threshold *d*_*t*_ which is reached for most bases on the genome. Using the average sequence depth we can also estimate the likelihood of having a certain number *v* of sequence variants in the pile observed. Reads *i* with a count *H*(i) close to or exceeding 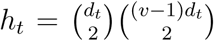 (we have fixed *r*_*1*_ to 1 and one of *r_4_, r_5_* or *r*_*6*_ to *r*) are most likely not in the same class at *R*_*j*_. Reads *i* in the same class as *R*_*j*_ should have a *H*(*i*) equal or close to zero.

## 3 Results

We have implemented both splitting methods. They are freely available as the programs split_agr and split_dis in the daccord package (see https://github.com/gtl/daccord). The daccord program (see [2]) in this package was also used to compute preliminary consensus sequences for the splitting. Read versus read alignments were computed using DALIGNER (cf. [1]). We performed two types of performance tests, both of which are based on simulated reads to ensure we can properly check whether and to what degree splittings computed are accurate.

In the first test we took a 190kb piece of the E. coli genome, duplicated it *k* times for *k* = 1, 2,… 8 and spiked in 1% difference between the duplicated versions and the original. The differences are single bp insertions, deletions and substitutions with equal probability. We generated reads of average length 15kbp with an error rate of 15% to evenly cover the sequences at depth *d* = 20. For the splitting we only considered read overlaps of 5kbp and more to reduce noise in the underlying statistics. Table 2 shows the performance of agreement and disagreement based splitting in this scenario. We provide precision (which fraction of the alignments kept is true), recall (which fraction of the true alignments is kept) and *F*_*1*_ (harmonic mean of precision and recall) score measures to quantify the performance of the read classification methods. All scores given are rounded to 3 significant decimals. While agreement based splitting has good precision for one and two modified copies, the performance quickly drops up to the point where essentially most wrong alignments are kept. The disagreement based splitting works close to perfectly in this setting. For computing the threshold *h*_*t*_ we have provided the correct value for the number of variants *v* to the program, as it does not yet support estimating it from the input data. The *δ* parameter was set to 1%.

**Table 2.** Performance of splitting on 190kbp stretch of E. coli with 1 — 8 copies added at 1% difference to original

|  | Agreement based |  |  | Disagreement based |  |  |
| --- | --- | --- | --- | --- | --- | --- |
| copies | Precision | Recall | $F_1$ score | Precision | Recall | $F_1$ score |
| 1 | 1 | 0.910 | 0.952 | 1 | 1 | 1 |
| 2 | 0.986 | 0.820 | 0.896 | 0.998 | 1 | 0.999 |
| 3 | 0.908 | 0.862 | 0.885 | 0.999 | 1 | 1 |
| 4 | 0.567 | 0.862 | 0.717 | 0.999 | 1 | 0.999 |
| 5 | 0.225 | 0.998 | 0.367 | 0.998 | 1 | 0.999 |
| 6 | 0.139 | 1 | 0.243 | 0.998 | 1 | 0.999 |
| 7 | 0.120 | 1 | 0.214 | 0.997 | 1 | 0.998 |
| 8 | 0.107 | 1 | 0.193 | 0.996 | 1 | 0.998 |

As the first test is highly synthetic, we have chosen a somewhat more realistic scenario for the second one. We have extracted regions containing the genes FCGR1(A |B|CP), FCGR2(A|B|C) and FCGR3(A|B) plus 100kbp to the left and right of these regions from chromosome 1 of the human reference genome (GRCh38). These regions are highly repetitive with repeating stretches of length up to 46kbp with a difference of merely 1% and one repeat of length 26kbp with 0.4% difference between the copies. We generated reads and alignments using the same parameters as for the other test. Table 3 shows the performance of the splitting approaches we measured. While the region considered is repetitive in it’s entirety, we do not have many cases of stretches appearing more than twice in total, i.e. most repeats have only two instances. As this is the setting in which agreement based splitting mostly works, we see a decent performance for this method as reflected in the table. For the disagreement based splitting we provide two lines, one for the default value of *h*_*t*_ = 441 which is computed as described above, the other one for *h_t_* = 7, the setting which maximises the *F*_1_ score in this scenario. As above we used *δ* = 1%. The recall value is good for both choices of *h_t_*. We lose hardly any true alignments. The precision value for the default *h*_*t*_ of 441 is worse than the one of the agreement based method. A closer look reveals that the average difference between the true sequences we fail to separate (which lead to the false positive alignments we keep) is 0.467% which is way below our setting for *δ*, so the failure in separation is not surprising. Just reducing the parameter *δ* below 1% however does not markedly improve the splitting, as this also greatly increases noise (disagreement tuples observed although they are not real). The solution to this may be to require longer (> 5kbp) overlaps between reads. As for PacBIO 50% of the sequenced bases is found in reads longer than 20kbp, this may be feasible. When we reduce *h_t_*, then we are able to rule out more false positives, as there are some tuples for which most of the disagreements are observed and not just the small fraction we assume as a lower bound in our statistical considerations. This however also reduces the recall.

**Table 3.** Performance of splitting on FCGR regions of human chromosome 1

| Agreement based |  |  | Disagreement based |  |  |  |
| --- | --- | --- | --- | --- | --- | --- |
| Precision | Recall | $F_1$ score | $h_t$ | Precision | Recall | $F_1$ score |
| 0.953 | 0.934 | 0.943 | 441 | 0.899 | 1 | 0.947 |
|  |  |  | 7 | 0.937 | 0.989 | 0.963 |

## 4 Conclusion

We have shown that repeat and haplotype separation in long reads with current read length and error rates is possible down to a difference of 1% and possibly less. This improves on the current state of the art of 3% set by Canu. The methods proposed also work if there are more than two underlying sequence versions. We hope these new insights can help to significantly improve the assembly of repetitive regions in genomes.

## Acknowledgments

We thank Gene Myers for interesting algorithmical discussions related to this paper.

